# Maturation signatures of conventional dendritic cell subtypes in COVID-19 reflect direct viral sensing

**DOI:** 10.1101/2021.03.03.433597

**Authors:** Laura Marongiu, Giulia Protti, Fabio A. Facchini, Mihai Valache, Francesca Mingozzi, Valeria Ranzani, Anna Rita Putignano, Lorenzo Salviati, Valeria Bevilacqua, Serena Curti, Mariacristina Crosti, Mariella D’Angiò, Laura Rachele Bettini, Andrea Biondi, Luca Nespoli, Nicolò Tamini, Nicola Clementi, Nicasio Mancini, Sergio Abrignani, Roberto Spreafico, Francesca Granucci

## Abstract

Growing evidence suggests that conventional dendritic cells (cDCs) undergo aberrant maturation in COVID-19 and this negatively affects T cell activation. The presence of functional effector T cells in mild patients and dysfunctional T cells in severely ill patients suggests that adequate T cell responses are needed to limit disease severity. Therefore, understanding how cDCs cope with SARS-CoV-2 infections can help elucidate the mechanism of generation of protective immune responses. Here, we report that cDC2 subtypes exhibit similar infection-induced gene signatures with the up-regulation of interferon-stimulated genes and IL-6 signaling pathways. The main difference observed between DC2s and DC3s is the up-regulation of anti-apoptotic genes in DC3s, which explains their accumulation during infection. Furthermore, comparing cDCs between severe and mild patients, we find in the former a profound down-regulation of genes encoding molecules involved in antigen presentation, such as major histocompatibility complex class II (MHCII) molecules, β_2_ microglobulin, TAP and costimulatory proteins, while an opposite trend is observed for proinflammatory molecules, such as complement and coagulation factors. Therefore, as the severity of the disease increases, cDC2s enhance their inflammatory properties and lose their main function, which is the antigen presentation capacity. In vitro, direct exposure of cDC2s to the virus recapitulates the type of activation observed in vivo. Our findings provide evidence that SARS-CoV-2 can interact directly with cDC2s and, by inducing the down-regulation of crucial molecules required for T cell activation, implements an efficient immune escape mechanism that correlates with disease severity.

## Results and Discussion

Clinical outcomes of COVID-19 are highly variable. Patients may show either no/mild symptoms (such as mild fever and cough) or severe respiratory involvement requiring hospitalization. In the most severe cases, Acute Respiratory Distress Syndrome (ARDS) can develop, with high levels of inflammatory hallmarks in the blood ^1, 2^ and diffuse intravascular coagulation (DIC) ^3,4^. In a non-negligible number of cases, COVID-19 is lethal ^5^. Patients presenting severe symptoms show immune dysregulation characterized by excessive release of type 1 and type 2 cytokines ^2^, and alterations of lymphoid and myeloid populations in the peripheral blood ^6^. Severe patients, diversely from mild patients, also show alterations in both Th17 and Th1 cell activation, with defects in the acquisition of effector functions ^7^.

Cells of myeloid origin play a pivotal role during infections by sensing pathogens, producing inflammatory mediators and by contributing to the activation of adaptive immunity. In this context, dendritic cells (DCs) are particularly relevant since they are specialized in antigen presentation and T cell priming ^8^. Given the functional specialization of DCs, the differences observed in the activated T cell compartments in severe versus mild patients suggest that alterations in activation may also be present in the conventional DC (cDC) compartment in patients presenting with different levels of disease severity.

cDCs have been divided in two subtypes, cDC1 and cDC2, originating from a common precursor (pre-DCs) ^9,10,11,12^, cDC1s have a high intrinsic capacity to cross-present antigens, due to the expression of the CLEC9A c-type lectin ^13^, and activate CD8^+^, Th1 and NK cells ^14^. Myeloid cDC2s express different Pattern Recognition Receptors (PRRs) and can promote a wide range of immune responses and especially CD4^+^ T cell responses ^15^. Recently, cDC2s have been divided in two subsets, DC2 and DC3 ^16,17,18^. DC3s are a heterogeneous population and comprehend non inflammatory cells showing a CD163^-^CD14^-^CD5^-^ phenotype, inflammatory CD163^+^CD14^+^CD5^-^ cells and a CD163^+^CD14^-^CD5^-^ intermediate subpopulation ^16^.

Functional impairments of cDCs have been described in COVID-19 patients, with decreased numbers in the blood ^19,20^, but not in bronchoalveolar lavage (BAL) samples ^7^, and reduced functionality, in terms of cytokines production and T cell priming capacity when restimulated in vitro ^21,22^. Nevertheless, a defect of maturation following in vitro restimulation does not necessarily indicate functional impairment, as activated DCs may not further respond to PRR agonists. Therefore, no specific information is available concerning the impact of SARS-CoV-2 infection on the maturation of DC subtypes and a better understanding is mandatory given the specific role of cDC subtypes in the activation and skewing of adaptive immune responses that ultimately contribute to COVID-19 pathogenesis ^23^.

In seeking for the impact of SARS-CoV-2 infection on blood DC subtypes, we analyzed peripheral blood DCs from severe and mild COVID-19 patients, according to World Health Organization (WHO) classification. Patients were enrolled from the STORM cohort (see **Supplementary Table 1** for the clinical data of the patients) of San Gerardo Hospital in Monza, Italy.

cDC1s were identified as CLEC9A^+^ and cDC2s as CD1c^+^FcεRIα^+^ over the CD11c^+^MHCII^+^ PBMCs excluding cells expressing markers for T and B lymphocytes (CD3 and CD19, respectively) and monocytes (CD88 and CD89) ^16^. CD14 was included in the analysis to identify DC3s ^17^ (**Supplementary Fig. S1 for gating strategies**). Consistent with some studies ^20^ but not others ^6^, we found a decreasing trend in the frequency of cDC1s and DC2s and an increasing trend in DC3s in COVID-19 patients compared with healthy donors (HDs) (**Fig. 1A**).

**Fig. 1.**
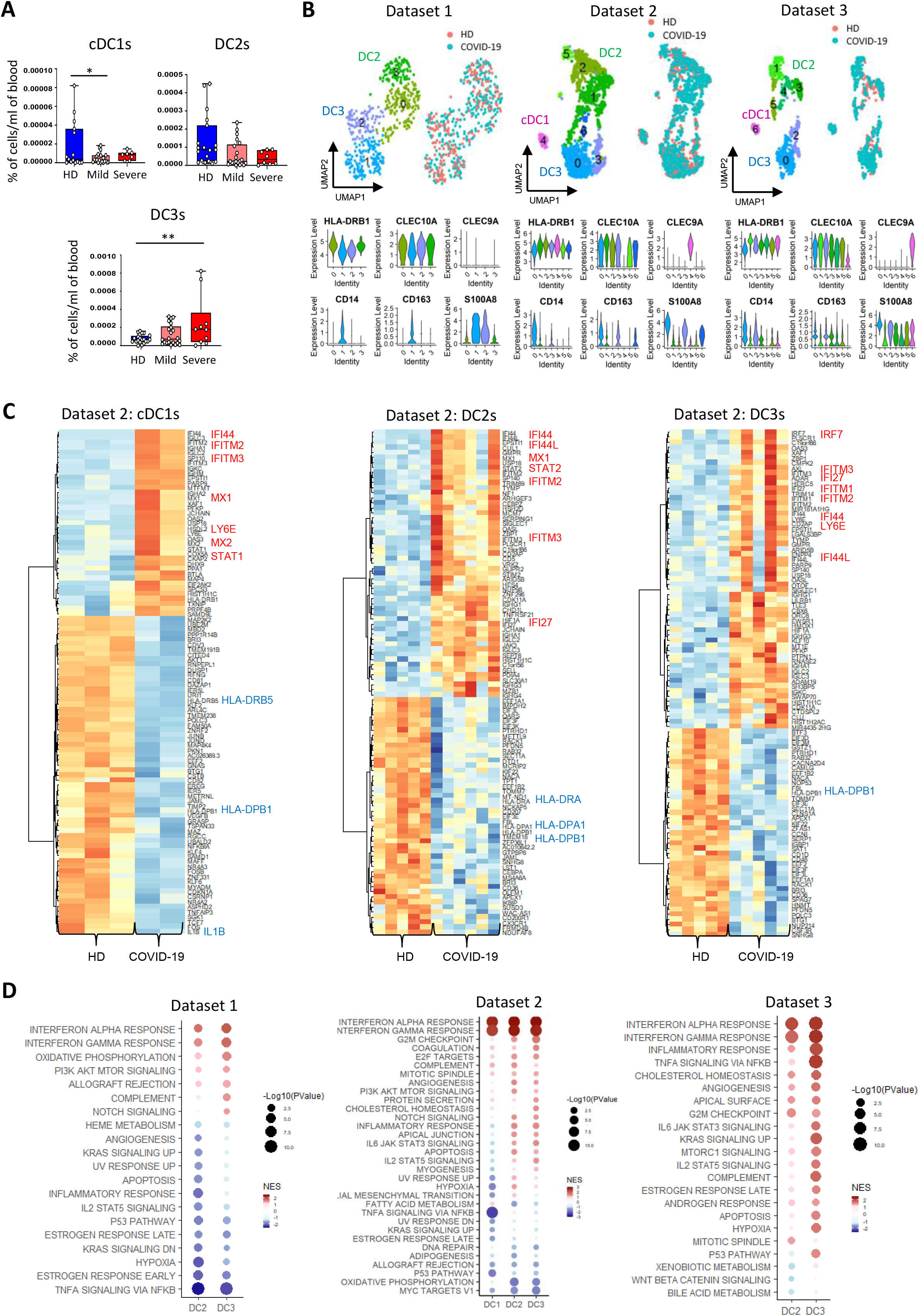
The response of cDCs to SARS-CoV-2 infection is dominated by ISGs. (A) Percentage of cDC1s, DC2s and DC3s on CD45^+^ cells from whole blood of COVID-19 patients (n=22 mild and n=10 severe) and HDs (n=21). Statistical significance was determined using one-way analysis of variance, followed by Sidak’s multiple comparison test. *p < 0.05; **p < 0.01. (B, upper panels) UMAP representations of cDC subtypes identified from the three scRNA-seq datasets analysed: dataset 1 (newly generated) and datasets 2 and 3 (publicly available). Cells are colored according to cDC subtype and donor origin (pink, HDs; lightblue, COVID-19). (B, lower panels) Violin plots illustrating expression levels of selected marker genes used for the manual annotation of cDC subtypes. (C) Heatmaps showing the top 100 DEGs for cDC1, DC2 and DC3 subsets comparing COVID-19 patients and HDs from dataset 2. Selected up-regulated genes (ISGs) are marked in red and down-regulated genes in blue. Ribosomal protein (RP) genes were removed from the top 100 DEGs. (D) GSEA of DEGs using the Hallmark collection: dataset 1 (left panel), dataset 2 (middle panel) and dataset 3 (right panel). For each DC subset, top 15 pathways based on significance are shown. NES, normalized enrichment score.

To perform a systematic characterization of the transcriptional response of DCs to SARS-CoV-2 infection, we analyzed three different single-cell transcriptomic datasets, two publicly available and a newly generated one. The analysis of three independent datasets allowed us to identify consistently altered signaling pathways, minimizing the effects of possible biases in the single datasets.

The new dataset (dataset 1) was generated using a droplet-based single cell platform (10X Chromium) and contains scRNA-seq data of CD11c^+^ MHC-II^+^ cells isolated from PBMCs of three COVID-19 patients (two mild and one severe) and two HDs (**Supplementary Table 1**). The second dataset ^22^ (dataset 2) contains cellular indexing of transcriptomes and epitopes by sequencing (CITE-seq) data of PBMCs and enriched DCs obtained from 7 COVID-19 patients (three mild and four severe) and 5 HDs, while the third dataset ^24^ (dataset 3) contains scRNA-seq data of PBMCs obtained from 18 COVID-19 patients (8 mild and 10 severe) and 21 HDs. Single-cell data from datasets 1 and 2 were first visualized using non-linear dimensionality reduction through uniform manifold approximation and projection (UMAP) and graph-based clustering algorithms (**Supplementary Fig. 2A, 3A**). Clusters containing myeloid DCs were identified based on the expression of markers that discriminate cDC2s and cDC1s from all other cell populations. Specifically, *CD1C*, *FCER1A* and *CLEC10A* were used to identify cDC2s, while *CLEC9A* was used to identify cDC1s (**Supplementary Fig. 2B, 3B**). For dataset 3, myeloid DCs already annotated by the authors were considered ^24^. Clusters corresponding to myeloid DCs in the three datasets were re-clustered in further iterations to separate cDC1s from cDC2s, to discriminate cDC2 subpopulations and to exclude possible contaminants. Specifically, DC3s were distinguished from DC2s based on the expression of *CD14*, *CD163* and *S100A8* markers. This approach allowed us to clearly identify cDC subsets (**Fig. 1B and Supplementary Fig. 3C,D**).

Next, in order to unravel the transcriptional response of each DC subset during SARS-CoV-2 infection, we aggregated cell-level counts into sample-level pseudobulk counts, mitigating single-cell mRNA measurement noise, and identified differentially expressed genes (DEGs) between COVID-19 patients and HDs (**Supplementary Table 2**). The low numbers of cDC1s allowed their analysis only in dataset 2.

In all DC subsets from the three datasets, when comparing expression profiles of COVID-19 patients with those of HDs, most of the genes up-regulated in COVID-19 were interferon (IFN) stimulated genes (ISGs) (**Fig. 1C and Supplementary Fig. 4A,B**). On the other hand, among the most significantly down-regulated genes in COVID-19, there were those encoding MHC class II molecules (**Fig. 1C**), indicating an impaired antigen presentation capacity of these cells. To understand more deeply which biological signaling pathways were differentially regulated in COVID-19 patients compared with HDs, we performed Gene Set Enrichment Analysis (GSEA) using two different gene sets: the Hallmark collection from the Molecular Signatures Database (MSigDB) and the literature-derived Blood Transcription Modules (BTMs) ^25^ (**Supplementary Table 3**). As it could be predicted by the identified DEGs, in all DC subtypes from the three datasets, maturation was dominated by ISGs while we could not detect the up-regulation of signatures containing classical activation markers and cytokines for T cell priming (**Fig. 1D and Supplementary Fig. 5A).** Together with the IFN-induced pathways, IL-6 pathways (IL-6-JAK-STAT3 and PI3K-AKT-mTOR ^26^) were recurrently up-regulated in cDC2s in all datasets (**Fig. 1D**). This is consistent with the relevance of IL-6 in COVID-19 pathogenesis and the expansion of activated Th17 cells in COVID-19 patients^27^.

The lack of a conventional maturation signature (lack of up-regulation of genes encoding MHCII and costimulatory molecules and cytokines) in circulating DCs prompted us to ask whether it was, in fact, possible to identify activated DCs in the blood. It could not be excluded that mature DCs reach circulation too late after activation when they are exhausted and, thus, only very late transcriptional events are visible.

We, therefore, investigated the transcriptional responses of circulating DC2 and DC3 subsets at single-cell resolution in different clinical conditions. Two distinct publicly available datasets were analyzed: the dataset from Reyes *et al*. ^28^ containing scRNA-seq data of PBMCs and enriched DCs obtained from patients with urinary tract bacterial infections of increasing severity (localized infection [Leuk-UTI], systemic infection with transient [Int-URO] or persistent organ dysfunctions [URO]), and the dataset from Hao *et al*. ^29^ containing CITE-seq data of PBMCs obtained from healthy volunteers that received an adenovirus-based vaccine. As previously described, we performed dimensionality reduction and unsupervised clustering to identify DC subpopulations. Our approach clearly identified DC subsets in both datasets (**Fig. 2A and Supplementary Fig. 6**). We then determined DEGs in infected or vaccinated donors with respect to the corresponding HDs (**Supplementary Table 4**) and performed GSEA.

**Fig. 2.**
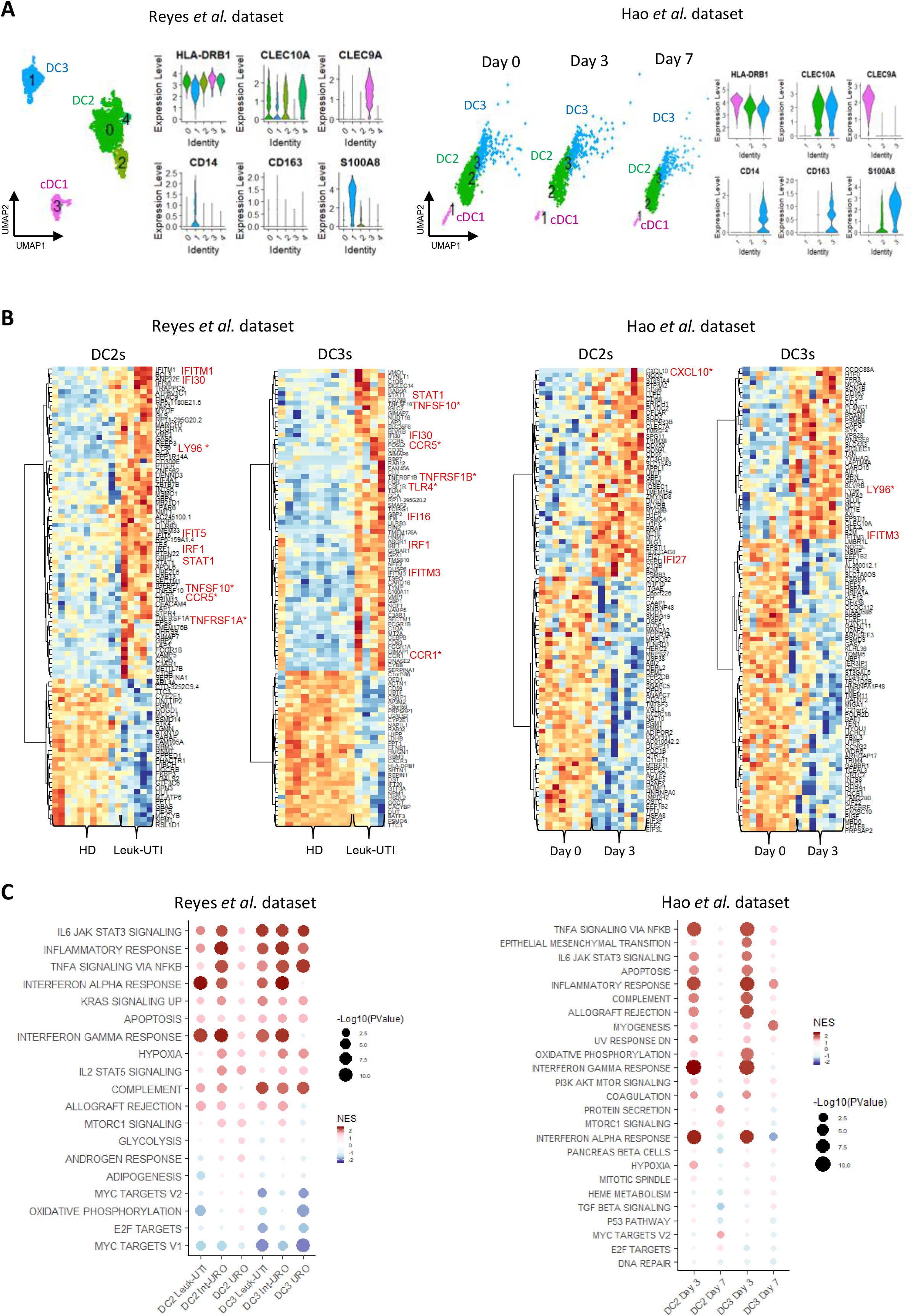
Activation signature of cDCs during bacterial infections and adenovirus-based vaccine administration. (A) UMAP representations of cDC subtypes and corresponding violin plots illustrating expression levels of selected marker genes used for the manual annotation of cDC subtypes: Reyes *et al*. dataset (left panel) and Hao *et al*. dataset (right panel). (B) Heatmaps showing the top 100 DEGs for DC2 and DC3 subsets comparing: Leuk-UTI patients with HDs from Reyes *et al*. dataset (left panel) and vaccinated donors at day 3 with unvaccinated donors from Hao *et al*. dataset (right panel). Selected up-regulated genes are marked in red. Asterisk indicates genes associated with pro-inflammatory functions. Ribosomal protein (RP) genes were removed from the top 100 DEGs. (C) GSEA of DEGs using the Hallmark collection: Reyes *et al*. dataset (left panel) and Hao *et al*. dataset (right panel). For each DC subset, top 10 pathways based on significance are shown. NES, normalized enrichment score. Leuk-UTI, urinary tract infection with leukocytosis. Int-URO, intermediate urosepsis. URO, urosepsis.

The results were in stark contrast to those obtained from COVID-19 patients. Indeed, in both datasets, circulating DC2s and DC3s showed an up-regulation not only of IFN pathways (as in COVID-19) but also of inflammatory signatures and genes relevant for immune responses (differently from COVID-19) (**Fig. 2B,C and Supplementary Fig. 7,8**). Among the most highly up-regulated genes there were several encoding activation molecules such as *CCR1*, *CCR5*, *CXCL10*, *TNFSF10* (*CD253/TRAIL*) as well as Toll-like receptor (TLR) genes, (**Fig. 2B and Supplementary Fig. 7A,B**).

These findings were confirmed by pathway analysis, which showed a clear up-regulation of activation pathways in DC2 and DC3 subsets in response to bacterial infections or vaccine, such as the inflammatory response pathway and the TNF-α signaling pathway (**Fig. 2C and Supplementary Fig. 8A,B**). Among the leading edge genes driving the enrichment of the inflammatory response pathway in response to bacterial infections there were several ones relevant for T cell activation (*IL1B*, *CCL5*, *TNFSF10*, *GPR183*, *CD69*, *SELL*) (**Supplementary Fig. 8C**).

Interestingly, we observed a stronger activation response of circulating cDC2s in patients with localized bacterial infections (Leuk-UTI group) and transient organ dysfunction (Int-URO group) than in patients with bacterial sepsis and persistent organ dysfunction (URO group) (**Fig. 2C**). This was expected since sepsis induces functional impairment of myeloid cells. These results indicate that conventionally activated DCs can be detectable in the blood when infections are both localized and systemic. Therefore, the lack of a classic activation signature, observed in cDC subtypes from blood of COVID-19 patients, is not a generalized phenomenon.

Recent studies have indicated potential functional differences between DC2 and DC3 subpopulations ^30^, in particular in inflammatory diseases, like Systemic Lupus Erythematosus (SLE), in which type I IFNs play a major role ^16^. In order to investigate a potential specific role for DC3s with respect to DC2s, we determined the genes differentially induced/downmodulated by these two subpopulations in response to SARS-CoV-2 stimulation and compared them to bacterial sepsis.

To increase our resolution, we pooled cDCs from the three COVID-19 datasets and performed Harmony integration ^31^, followed by graph-based clustering. After integration, we obtained 2,415 cDCs (**Fig. 3A**) and we clearly identified clusters of cDC1s, DC2s and DC3s (**Fig. 3B**). Only 24 genes (p-value < 0.05 and absolute log2FC > 1) were differentially expressed in DC3s compared with DC2s in response to COVID-19 infection, of which 17 were up-regulated and 7 were down-regulated (**Fig. 3C, left panel** and **Supplementary Table 5**). On the other hand, 152 genes (p-value < 0.05 and absolute log2FC > 1) were identified as differentially regulated in DC3s compared with DC2s in response to intermediate urosepsis (Int-URO condition), of which 59 were up-regulated and 93 were down-regulated (**Fig. 3C, right panel** and **Supplementary Table 5**).

**Fig. 3.**
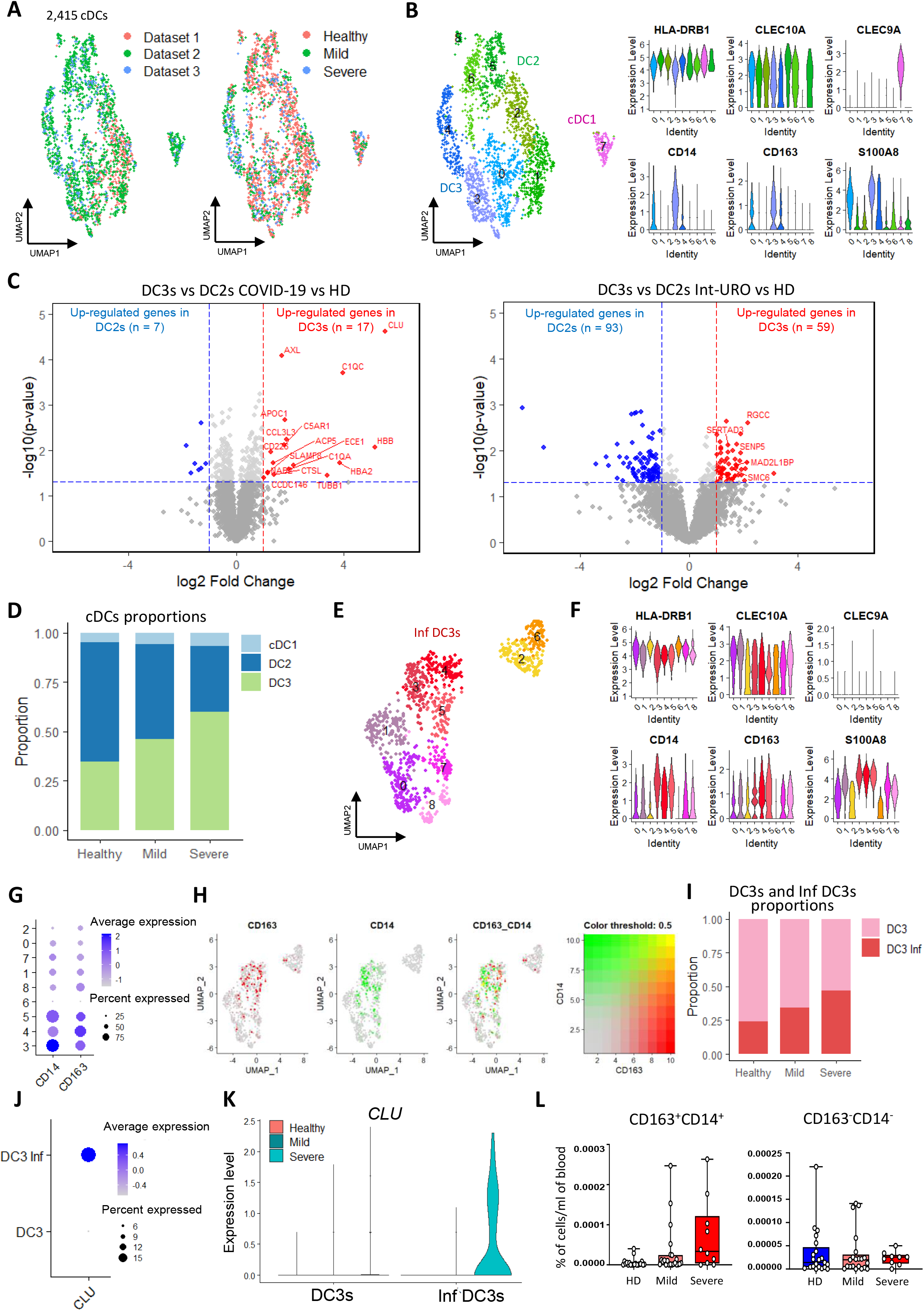
DC2s and DC3s respond similarly to SARS-CoV-2 infection and inflammatory DC3s accumulate in severe patients. (A) UMAP representations of cDCs after datasets integration. Cells are colored according to dataset origin (left panel; pink, dataset 1; green, dataset 2; lightblue, dataset 3) and clinical condition (right panel; pink, HDs; green, mild patients; lightblue, severe patients). (B) UMAP representation of cDC subtypes and corresponding violin plots illustrating expression levels of selected marker genes used for the manual annotation of cDC subtypes. (C) Volcano plots showing genes differentially induced in DC3s compared with DC2s in response to COVID-19 (left panel) and intermediate urosepsis (Int-URO condition, Reyes *et al*. dataset, right panel). Genes with p-value < 0.05 and absolute log2 fold change > 1 were considered significant. Selected genes are highlighted (red: up in DC3s; blue: up in DC2s). (D) Barplots showing the relative abundance of cDC populations in HDs, mild and severe patients. (E) Subclustering of DC3 population to identify inflammatory DC3s (clusters 3, 4 and 5). (F-H) Expression levels of selected marker genes used for the identification of inflammatory DC3s within the DC3 population: (F) violin plots showing expression levels of *HLA-DRB1*, *CLEC10A*, *CLEC9A*, *CD14*, *CD163* and *S100A8*, (G) dotplot showing expression levels of *CD14* and *CD163* across DC3 clusters and (H) combined feature plot demonstrating co-expression of CD14 and CD163 in clusters 3, 4 and 5. (I) Barplots showing the relative abundance of DC3s and inflammatory DC3s in HDs, mild and severe patients. (J-K) Expression level of the anti-apoptotic gene *CLU* in DC3s and inflammatory DC3s is shown as (J) dotplot and (K) split violin plot by clinical condition (pink, HDs; blu, mild patients; lightblue, severe patients). (L) Percentage of inflammatory (CD14^+^CD163^+^) and non-inflammatory (CD14^-^CD163^-^) DC3s on CD45^+^ cells from whole blood of COVID-19 patients (n=22 mild and n=10 severe) and HDs (n=21).

The diversity in the responses of DC3s and DC2s during bacterial infections could be, at least partly, explained by the differential expression of some receptors, such as CD14 exclusively expressed by DC3s. CD14 is a component of the receptor complex of lipopolysaccharide (LPS), a major factor of the outer membrane of Gram-negative bacteria, and contributes to LPS recognition and internalization of the receptor complex ^32^. Therefore, thanks to the expression of CD14, DC3s can respond more efficiently to Gram-negative bacteria than DC2s. CD14 has also important roles as chaperon for ligands of endosomal and cytosolic PRRs^32^. Therefore, the differences between DC2 and DC3 responses observed in Gram-negative bacterial infections may also occur after Gram-positive bacterial recognition.

In conclusion, these findings suggest that DC2s and DC3s respond in a very similar way to SARS-CoV-2 infections, while they show more diversified responses to bacterial infections. Interestingly, among the few genes differentially expressed between DC3s and DC2s in response to COVID-19, there were genes encoding complement factors and receptors (*C1QC*, *C1QA* and *C5AR1*) and, most importantly, anti-apoptotic genes such as *AXL* and *CLU* that resulted the most significantly up-regulated (**Fig. 3C, left panel)**. This suggests that DC3s are less susceptible to apoptosis than DC2s and may explain why they tend to increase while all other cDC populations decrease during SARS-CoV-2 infection (**Fig. 1A**). When comparing these results with those obtained from bacterial infections, we found that genes associated with cell cycle progression and cell proliferation (*RGCC*, *SENP5*, *SMC6*, *SERTAD3*, *MAD2L1BP*) were specifically up-regulated in DC3s (**Fig. 3C, right panel**). Therefore, DC3s may proliferate during inflammatory responses or circulating DC3s may contain some proliferating progenitors that expand the DC3 population during bacterial infections.

Altogether, these observations could explain why DC3s increase in number in acute and chronic inflammatory conditions ^16,33^. Moreover, the higher persistence potential of DC3s induced by inflammation could explain why their frequency is highly variable in HDs.

Coherently with these observations, we found an alteration of cDCs relative abundance in COVID-19 patients compared with HDs also at single-cell resolution. Specifically, DC3s showed increased frequencies in patients, which positively correlated with disease severity (**Fig. 3D**). Since DC3s are highly heterogeneous, we analyzed DC3 subpopulations at higher resolution. Hence, we retained clusters corresponding to DC3s and performed a re-clustering procedure to discriminate between DC3s and inflammatory DC3s (**Fig. 3E**). Interestingly, we identified three main phenotypes, characterized by low expression of both CD14 and CD163 (clusters 2 and 6), intermediate expression (clusters 0, 1, 7 and 8) and high expression (clusters 3, 4, 5), reflecting the heterogeneity of DC3 population (**Fig. 3F,G,H**). Clusters with the highest expression of both CD14 and CD163 (clusters 3, 4, 5) were annotated as inflammatory DC3s. We found a progressive increase in the relative abundance of inflammatory DC3s from HDs, to mild and finally to severe patients (**Fig. 3I**). This result was associated with the higher expression of the anti-apoptotic gene *CLU* in the inflammatory DC3s population, specifically in critically ill patients (**Fig. 3J,K**). Accordingly, an increasing trend in the frequency of inflammatory DC3s (CD14^+^CD163^+^), and not of non-inflammatory DC3s (CD14^-^CD163^-^), was observed in the blood of severe patients (**Fig. 3L**).

In order to seek for specific alterations in the innate immune signature of mild and severe COVID-19 patients and to link immune response variation to disease severity, we investigated cDC2 gene expression profiles in severe versus mild COVID-19 patients. As previously described, we aggregated cell-level counts into sample-level pseudo-bulk counts and identified DEGs between severe and mild COVID-19 patients (**Supplementary Table 6**).

In both DC2s and DC3s, we identified an important number of DEGs (200 for DC2s and 169 for DC3s, p-value < 0.05 and absolute log2FC > 1) between severe and mild patients, indicating relevant differences in the transcriptional response of these two groups (**Fig. 4A**). Interestingly, inflammatory genes not directly related to the activation of adaptive immunity, like complement factors (*C1QC, C1QB*) and complement receptors (*C5AR1*), genes involved in the production of leukotrienes known to exacerbate respiratory syndromes (*ALOX5AP*), genes of the coagulation cascade (*THBS1*, *THBD*), factors involved in vasodilation (*ADM*) and other inflammatory genes like *CD14*, *S100A8/A9*, *ADAM9* and *CD163* were significantly up-regulated in severe versus mild patients in DC2s and/or DC3s (**Fig. 4A**). In addition, genes that negatively interfere with the maturation of DCs finalized to T cell activation, like *TMEM176A*, *CD109*, *MT1E* were up-regulated in severe compared to mild patients (**Fig. 4A**). Moreover, further supporting what discussed above, the anti-apoptotic gene *CLU* was found to be among the most significantly up-regulated genes in DC3s of severe patients (**Fig.4A, right panel**).

**Fig. 4.**
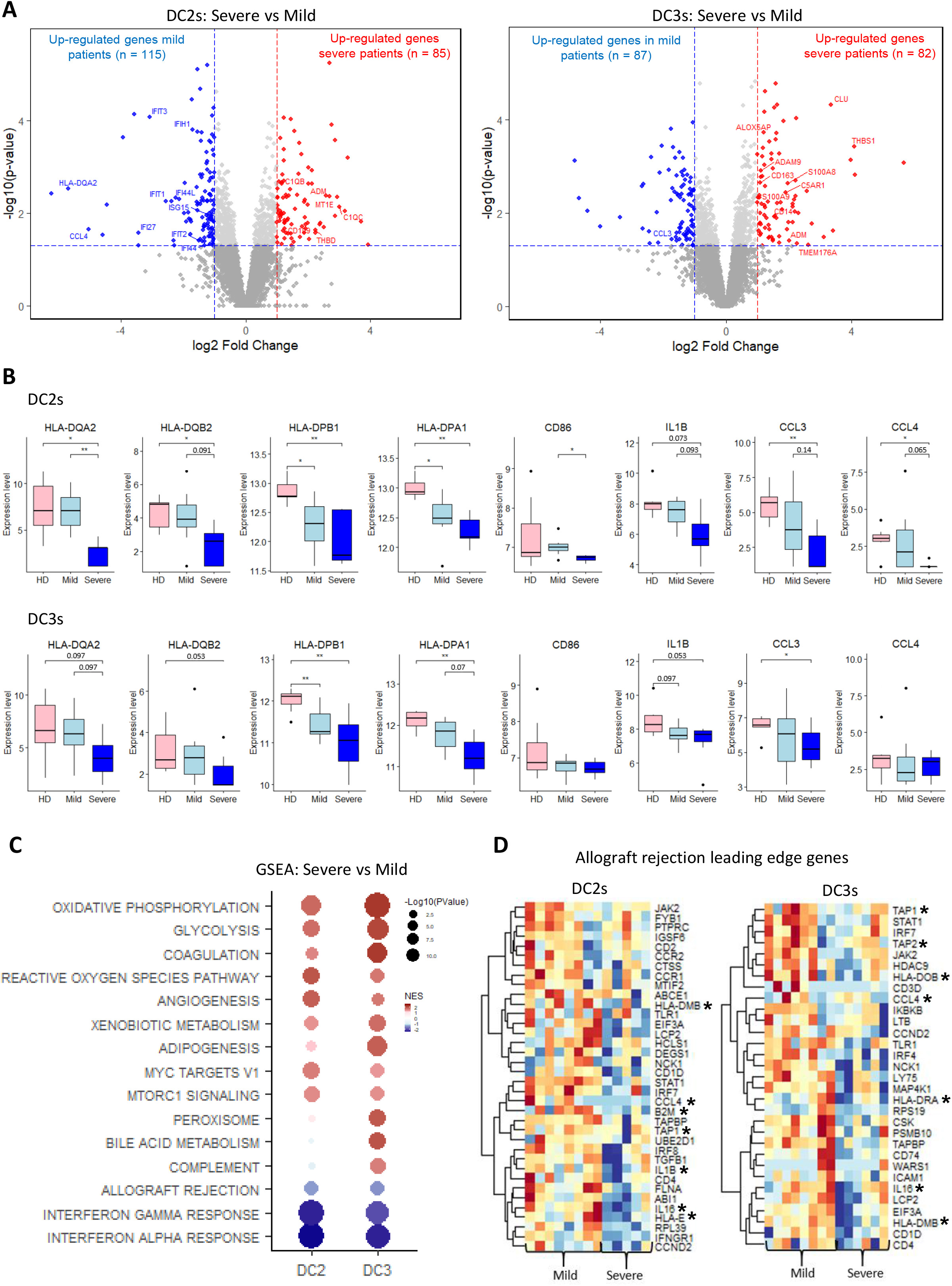
cDC2s enhance their inflammatory properties and lose antigen presentation capacity in severe COVID-19 patients. (A) Volcano plots showing genes differentially expressed in severe compared with mild patients in DC2s (left panel) and DC3s (right panel). Genes with p-value < 0.05 and absolute log2 fold change > 1 were considered significant. Selected genes are highlighted (red: up in severe patients; blue: up in mild patients). (B) Boxplots showing expression levels of selected genes in DC2s and DC3s in HDs, mild and severe patients. Statistical analyses were performed using Wilcoxon rank sum test. * p < 0.05; ** p < 0.01. (C) GSEA of DEGs in severe compared with mild patients using the Hallmark collection. For each DC subset, top 10 pathways based on significance are shown. NES, normalized enrichment score. (D) Heatmaps showing leading edge genes of the allograft rejection pathway in mild and severe patients. Asterisk indicates genes associated with fundamental functions for DC-mediated T cell activation.

Strikingly, genes encoding MHCII molecules, the costimulatory molecule CD86 and cytokines, such as *IL1B, CCL3* and *CCL4*, showed a progressive down-regulation from HDs to mild and finally severe patients (**Fig. 4B**).

These observations were consolidated by pathway analysis, which showed a clear up-regulation of pathways involved in metabolism, coagulation, angiogenesis and reactive oxygen species in severe compared with mild patients, and a down-regulation of IFN pathways (**Fig. 4C**). Interestingly, among the leading edge genes of the allograft rejection pathway, that was found to be down-regulated in severe compared with mild patients, there were many genes critical for DC-mediated T cell activation, such as those coding for proteins involved in antigen presentation on both MHCI and MHCII pathways (*B2M*, *TAP1*, *TAP2*, *HLA-DMB*, *HLA-DRA*) and genes encoding molecules relevant for T cell recruitment and activation (*IL16*, *IL1B*, *CCL4*) (**Fig. 4D**). Specific down-regulation of these genes in severely ill patients emphasizes the alteration of cDC functions in these individuals, which may be associated with a worse disease progression.

Altogether, these observations indicate that, as disease severity increases, cDC2s progressively skew toward inflammatory activities and lose the antigen presenting function. This could explain the alteration of the activated T cell compartment observed in severely ill patients.

Results shown till now indicate that, during COVID-19 infections, DC3s and DC2s respond similarly to the virus with three main features: i) the up-regulation of ISGs and IL-6 pathways; ii) a progressive down-regulation from mild to severe patients of genes encoding signal 1 and signal 2 molecules associated with antigen presentation; iii) the up-regulation of an inflammatory signature, mainly represented by complement and coagulation factors, in severe patients.

We wondered whether these features could be due to the exposure to mediators released during SARS-CoV-2 infection or to the direct interaction of cDCs with the virus. The first cDC characteristic observed in our study is compatible with both the direct interaction with the virus and the exposure to paracrine cytokines, such as IFNs and IL-6 produced by bystander cells. The lack of expression of IFN and IL-6 genes in circulating DCs does not necessarily mean that cDCs cannot be a source of these cytokines, since the expression of genes encoding these molecules is acutely regulated and may be shut down when DCs reach the circulation. By contrast, the systematic down-regulation of genes encoding MHCII molecules is more likely explained by a direct interaction of cDCs with the virus. This prediction was also supported by evidence that the virus can directly activate monocyte derived DCs following abortive infection ^34^.

Therefore, we investigated whether the direct interaction of cDC2s with the virus could induce a similar response to that observed at single-cell resolution. By using IL-6 and MHCII as readouts, we measured the response to the virus of cDC2s (CD1c^+^CD19^-^ cells) freshly isolated from HDs. As predicted, we found that SARS-CoV-2 directly induced a significant down-regulation of MHCII surface expression and the up-regulation of IL-6 in both DC2s and DC3s (**Fig. 5A,B)**. Diversely, the exposure of cDC2s to sera from mild and severe patients, that contain inflammatory cytokines and other mediators, could not induce any modification in MHCII and IL-6 expression (**Fig. 5C**). This suggests that at least part of the peculiar response of cDCs induced in vivo by SARS-CoV-2 infection can be directly imposed by the virus.

**Fig. 5.**
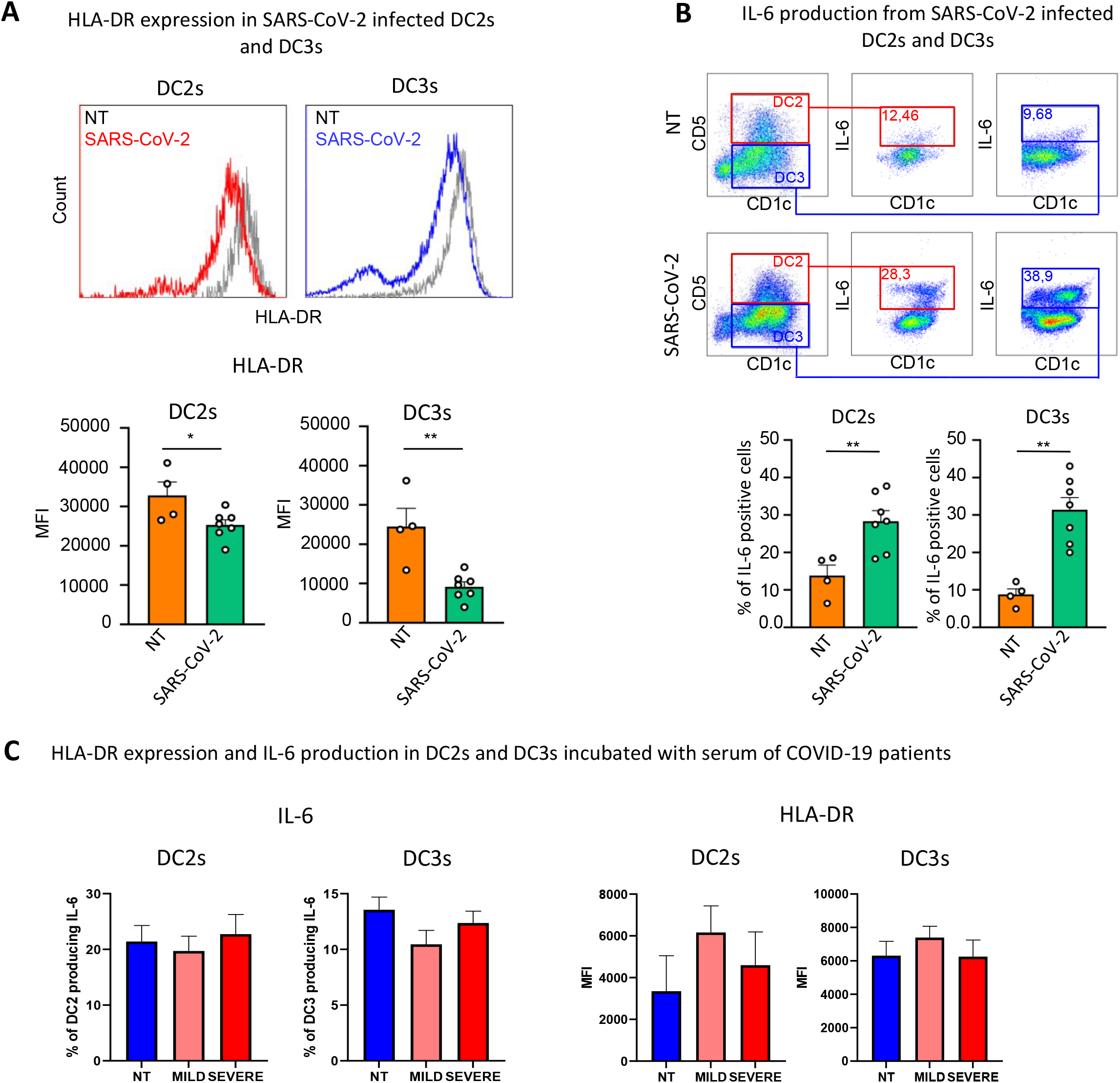
SARS-CoV-2 directly induces down-regulation of HLA-DR and production of IL-6 in DC2s and DC3s. (A, upper panel) Representative histograms showing HLA-DR expression in cDC2s from HDs infected or not (NT) with 0.4 MOI of SARS-CoV-2 for 18 hours. DC2s and DC3s were identified as CD5^+^ CD1c^+^ and CD5^-^ CD1c^+^ respectively over the CD11^+^LIN^-^ (CD88, CD89, CD3 and CD19) and FcεRIα^+^. (A, lower panel) Quantitative analysis of mean fluorescence intensity (MFI) of HLA-DR in DC2s and DC3s. Statistical significance was determined with unpaired student’s t-test. *p < 0.05, **p < 0.01; n=4 NT donors and n=7 donors for SARS-CoV-2 infection. (B, upper panel) Representative dot plots showing the percentage of IL-6 producing DC2s and DC3s after viral infection as described in B. (B, lower panel) Quantitative analysis of the percentage of IL-6 producing cells. Statistical significance was determined with unpaired student’s t-test. **p < 0.01; n=4 NT donors and n=7 donors for SARS-CoV-2 infection. (C, left panel) Quantitative analysis of the percentage of IL-6 producing DC2s and DC3s after 18h incubation with sera from n=4 mild and n=4 severe COVID-19 patients, NT (not treated). (C, right panel) Quantitative analysis of mean fluorescence intensity (MFI) of HLA-DR in DC2s and DC3s treated or not (NT) for 18h with sera from n=4 mild and n=4 severe COVID-19 patients.

In conclusion, SARS-CoV-2 can be detected directly by DCs and induces down-regulation of signals necessary for activation of T lymphocytes, a phenomenon that is accentuated with disease severity. This allows the virus to evade control of the adaptive immune system, while the host attempts to counteract viral infection with innate immunity.

Understanding how DCs manage SARS-CoV-2 infection will help identify ad hoc interventions to achieve optimal adaptive responses, a prerequisite for a good prognosis ^35,23^.

## Materials and Methods

### Flow cytometric analysis

PBMCs from COVID-19 patients enrolled from the STORM cohort were extracted from peripheral blood by density gradient centrifugation using Ficoll (GE Healthcare). Cells were washed twice and stained for 30 minutes on ice using the following anti-human antibodies (1:200, Becton Dickinson): anti-FcεRIα PE-Cy7, anti-CD14 PE, anti-CD1c APC-Cy7, anti-Clec9 (CD370) Alexa 647, anti-CD5 BV786, anti-CD3 BV605, anti-CD19 BV605, anti-CD88 BV605, anti-CD89 BV605, anti-CD11c BV480, anti-CD163 BV421, anti-HLA-DR BUV805. Cells were then washed and fixed using fixation buffer (Becton Dickinson) and acquired using BD FACSsymphony instrument (Becton Dickinson). Analyses were performed with Flow jo X software.

### cDC2s purification and activation

Human cDC2 cells were purified from peripheral blood mononuclear cells (PBMCs) extracts from buffy coat of healthy donors (provided by Niguarda hospital blood bank) by Ficoll-Paque density gradient centrifugation. Briefly, blood was stratified on Ficoll-Paque PLUS (GE Healthcare) in 3:4 ratio and centrifuged at 1500 r.p.m. for 30 min without brake. PBMCs were washed twice, collected and CD1c^+^ cells were purified using MACS beads according to the manufacturer’s instructions (Miltenyi Biotec). Cells were cultured in Roswell Park Memorial Institute (RPMI) 1640 medium (Euroclone) containing 10% heat-inactivated fetal bovine serum (Euroclone), 100 IU of penicillin, streptomycin (100 μg/ml), 2 mM l-glutamine (Euroclone). cDC2s were infected with 0.4 MOI of SARS-CoV-2 for 18h or treated with serum of COVID-19 patients (ratio serum/medium 1:1), then collected and stained with anti-FcεRIα PE-Cy7, anti-CD14 PE, anti-CD1c APC-Cy7, anti-CD5 BV786, anti-CD3 BV605, anti-CD19 BV605, anti-CD88 BV605, anti-CD89 BV605, anti-CD11c BV480, anti-CD163 BV421, anti-HLA-DR BUV805 (1:200, all from Becton Dickinson). Cells were then fixed and permeabilized with cytofix/cytoperm reagent kit (Becton Dickinson) and stained with anti-IL-6 FITC antibody, according to the manufacturer’s instructions. Samples were acquired with the BD FACSsymphony instrument (Becton Dickinson) and analyzed with Kaluza software.

### Single-cell RNA sequencing datasets analyzed in the study

In this study, three different single-cell datasets from COVID-19 patients and healthy controls were analyzed.

Dataset 1 was newly generated. Myeloid cells were sorted (CD11c^+^ MHCII^+^) from 3 COVID-19 patients (2 mild and 1 severe, enrolled from the STORM cohort) and 2 healthy donors using a MACSQuant Tyto (Miltenyi) (**Supplementary Fig. 1B**). After sorting, cell number and viability were evaluated using an automated cell counter. Viability for each sample was ≥75%. 10,000 cells per sample were loaded on a Chromium Next GEM Chip G (10x Genomics). A Chromium controller (10x Genomics, Pleasanton, CA, USA) was used to generate single-cell GEMs, according to Chromium Next GEM Single Cell 5’ Library & Gel Bead Kit v1.1 protocol (PN-1000165; 10x Genomics). Full-length cDNA amplification and 5’ gene expression library construction were performed according to manufacturers’ instructions in a Veriti 96-well Thermal Cycler (Thermo Fisher Scientific). Indexed libraries were sequenced on an Illumina Novaseq 6000 platform, on a S2 flowcell, 150bp PE (20,000 read pairs per cell). Reads from FASTQ files were aligned against the GRCh38 human reference genome and quantified using the Cell Ranger pipeline (10x Genomics) version 3.0 with default parameters. Single cell data have been deposited in GEO (GSE168388) and will be accessible upon publication.

Dataset 2 ^22^ is a CITE-seq experiment with PBMCs and enriched DCs from 7 COVID-19 patients (three mild and four severe) and 5 HDs. Count matrices were downloaded from the Gene Expression Omnibus (GEO) (GSE155673).

Dataset 3 ^24^ is a scRNA-seq experiment with PBMCs from 18 COVID-19 patients (8 mild and 10 severe) and 21 HDs. Seurat objects were downloaded from FASTGenomics (https://www.fastgenomics.org/). Only cells annotated as myeloid DCs by the authors were used in downstream analyses.

To compare transcriptional responses of cDC subsets between SARS-CoV-2 infection and other inflammatory conditions, we analyzed two additional publicly available datasets.

The Reyes *et al*. dataset ^28^ is a scRNA-seq experiment with PBMCs and enriched DCs obtained from patients with bacterial infections and healthy controls. Briefly, subjects were enrolled in two different cohorts. A primary cohort contains subjects that were classified into three clinical categories: Leuk-UTI, Int-URO and URO. The Leuk-UTI group refers to subjects with urinary-tract infection (UTI) with leukocytosis (blood WBC ≥ 12,000 per mm^3^) but no organ dysfunction. The Int-URO (intermediate urosepsis) group contains subjects with UTI with mild or transient organ dysfunction, and the URO (urosepsis) group refers to subjects with UTI with clear or persistent organ dysfunction. Ten subjects were classified as Leuk-UTI, seven as Int-URO and ten as URO. A second cohort comprises hospitalized subjects classified into three conditions: subjects with bacteremia and sepsis not requiring intensive care unit (ICU) admission (Bac-SEP group, four subjects), subjects with sepsis requiring ICU care (ICU-SEP, eight subjects) and subjects in the ICU for conditions other than sepsis (ICU-NoSEP, seven subjects). Data were downloaded from the Broad Institute Single Cell Portal (https://singlecell.broadinstitute.org/single_cell) (SCP548). For downstream analysis, we retained monocytes and DCs as annotated by the authors.

The Hao *et al*. dataset ^29^ is a CITE-seq experiment with PBMCs from 8 healthy volunteers enrolled in an adenovirus-based HIV vaccine trial. For each subject, PBMCs were collected at three time points: immediately before (day 0), three days, and seven days following vaccine administration. Data were downloaded from https://atlas.fredhutch.org/nygc/multimodal-pbmc/. For downstream analysis, we retained only cDCs as annotated by the authors.

### Single-cell data processing and analysis

Data processing and analysis for all single-cell datasets was performed using the Seurat package (version 4.0) ^29^ in R (version 4.0.3).

First, filters were applied to remove low-quality cells. These were based on the number of genes and UMIs detected in each cell and on the percentage of reads mapping to mitochondrial genes (cells with < 500 genes and > 10% of reads mapping to mitochondrial RNA were removed). Counts were then normalized and log-transformed using sctransform ^36^, while regressing out UMI counts and percentage of mitochondrial counts.

For dimensionality reduction, PCA was performed. Principal components (PCs) were fed to Harmony ^31^ for batch correction and/or integration of datasets from both disease and healthy conditions. UMAP was used for 2D visualization. Clusters were identified with the shared nearest neighbour (SNN) modularity optimization-based clustering algorithm followed by Louvain community detection. Cell type assignment was manually performed using marker genes, as detailed in figures. cDCs were retained and re-clustered again to identify subsets.

### Pseudobulk differential gene expression analysis

After the identification of cDC subsets, we aggregated cell-level counts into sample-level pseudobulk counts. For each DC subset, only donors with at least 10 cells were retained.

For the dataset from Reyes *et al*., only samples from the primary cohort were considered for differential analysis due to the low number of DCs obtained from subjects from the secondary cohort.

Differential expression analysis was performed using the quasi-likelihood framework of the edgeR package ^37^, using each donor as the unit of independent replication.

### Gene Set Enrichment Analysis

Pre-ranked GSEA ^38^ was performed on the differentially expressed genes (DEGs) using the fgsea package ^39^. The Hallmark gene sets and the Blood Transcription Modules (BTM) ^25^ were used. BTM families analyzed in this study are reported in **Supplementary Table 3**.

### Integration between datasets

cDCs identified in the three COVID-19 datasets were pooled, integrated using Harmony ^31^, and further subclustered using the shared nearest neighbour (SNN) modularity optimization-based clustering algorithm followed by Louvain community detection with a resolution of 0.6 to identify cDC1, DC2 and DC3 clusters.

## Code availability

Code used for data analysis will be made available upon publication.

## Acknowledgments

RS thanks the QCB Collaboratory community directed by Matteo Pellegrini.

## Author contributions

Conceptualization: FG, LM

Methodology: LM, RS, GP, VR, ARP, LS, VB, CS, MCC, SA

Investigation: LM, GP, RS, FAF, MV, FM, SA, SC, MDA, LB, AB, LN, NT

Visualization: FG, GP, LM

Funding acquisition: FG

Supervision: FG

Writing – original draft: FG, GP

Writing – review & editing: FG, GP, LM

## Data and materials availability

All data are available in the main text or in the supplementary materials.

## Supplementary Figures and Tables

**Supplementary Figure 1.**
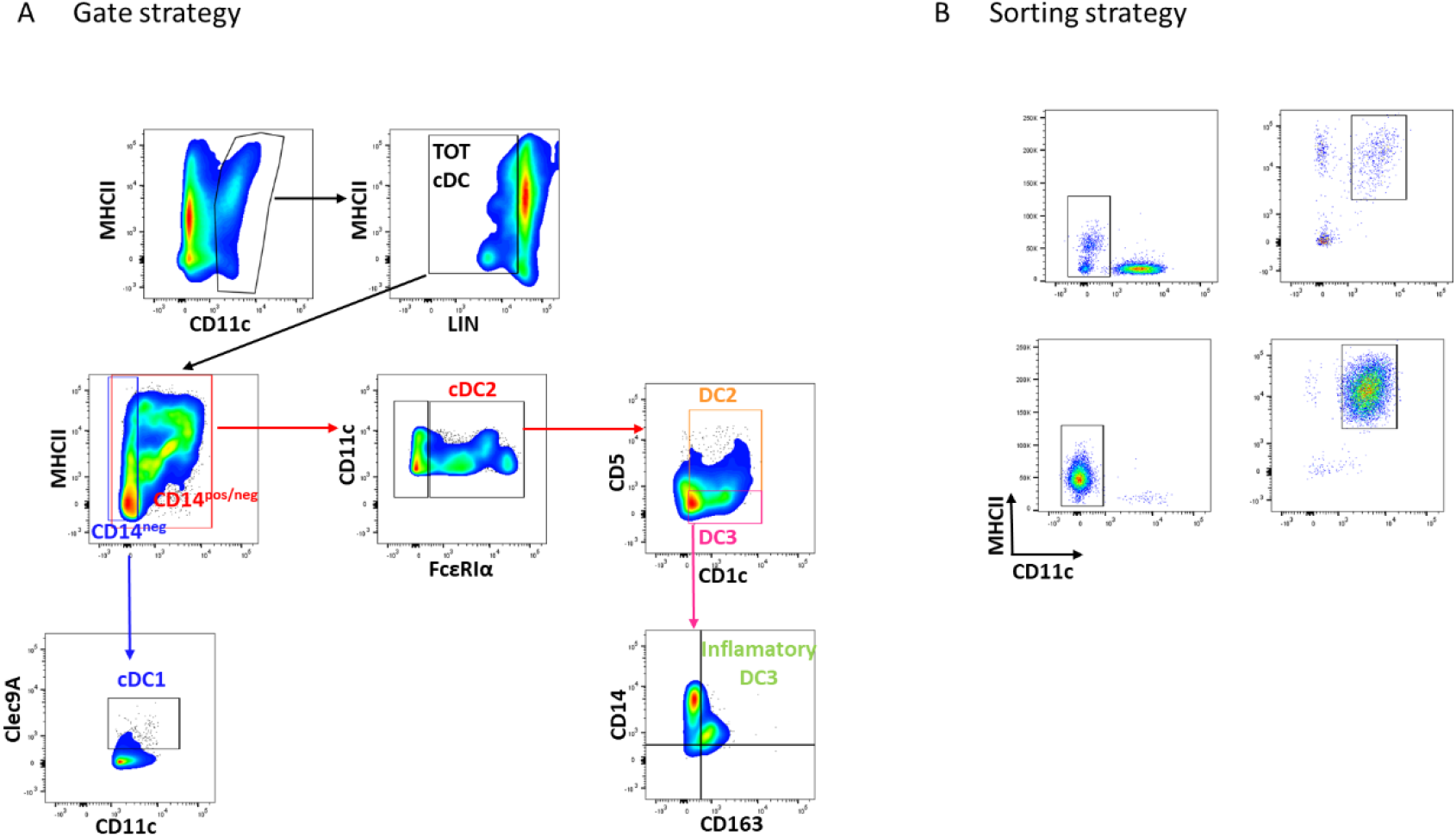
(A) Gating strategy to identify DCs subsets from PBMCs. Total DCs (cDCs TOT) were detected among the CD11c^+^ MHC-II^+^ and LIN^-^ (CD88, CD89, CD3 and CD19) population. cDC1s were identified as CLEC9A^+^ from the CD14^-^ fraction of total DCs. cDC2s (FcεRIα^+^) include CD14^+^ and CD14^-^ cells. DC2s and DC3s were identified as CD5^+^ CD1c^+^ and CD5^-^ CD1c^+^ respectively. Inflammatory DC3s were recognized as CD14^+^CD163^+^ cells.

**Supplementary Figure 2.**
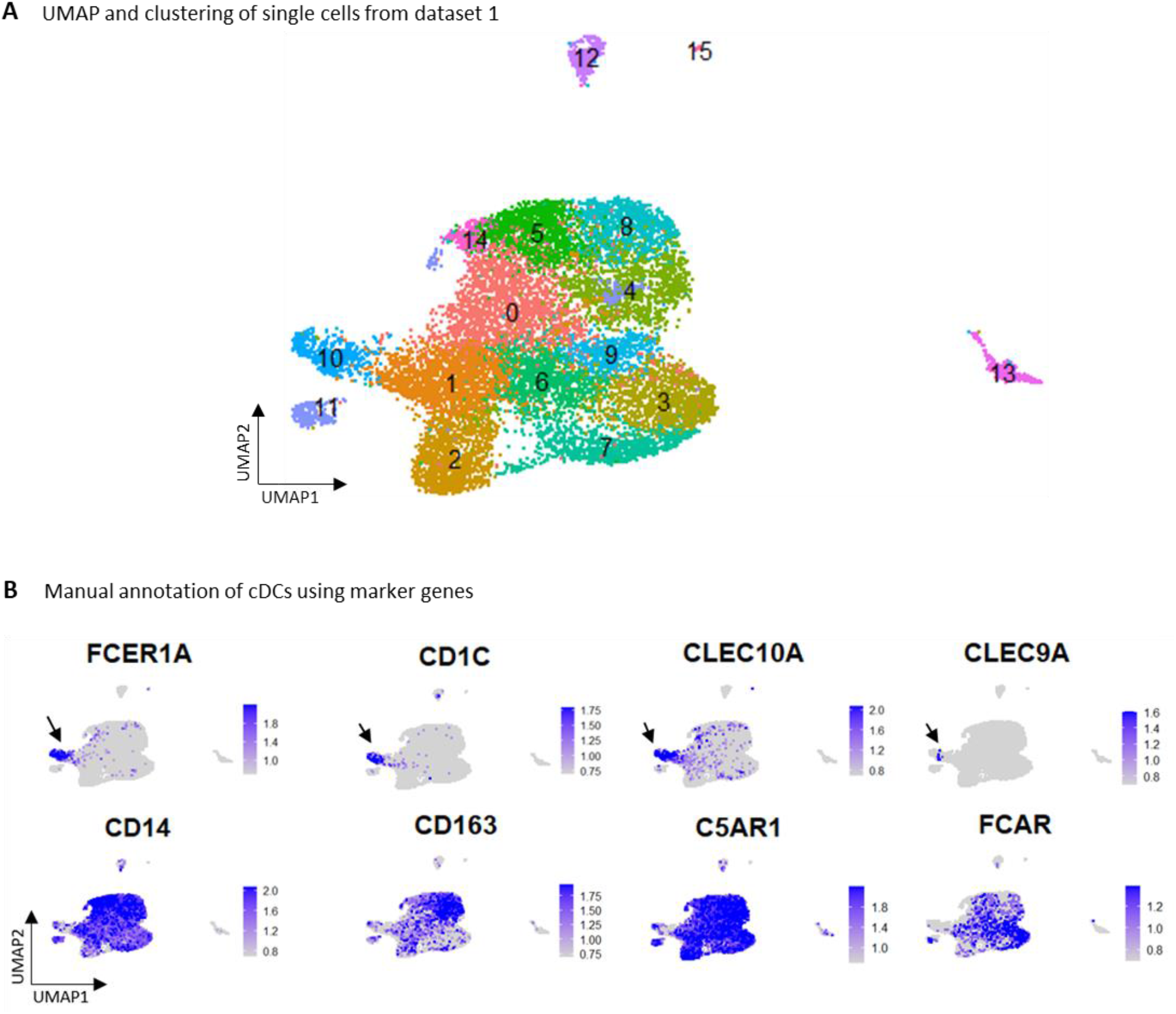
(A) UMAP and clustering of single cells from dataset 1. (B) Feature plots showing the expression levels of selected marker genes used to identify cDC cluster. Black arrows indicate cDC cluster (cluster 10). This cluster was re-clustered in a final iteration to clearly delineate cDC subsets as shown in Figure 1B.

**Supplementary Figure 3.**
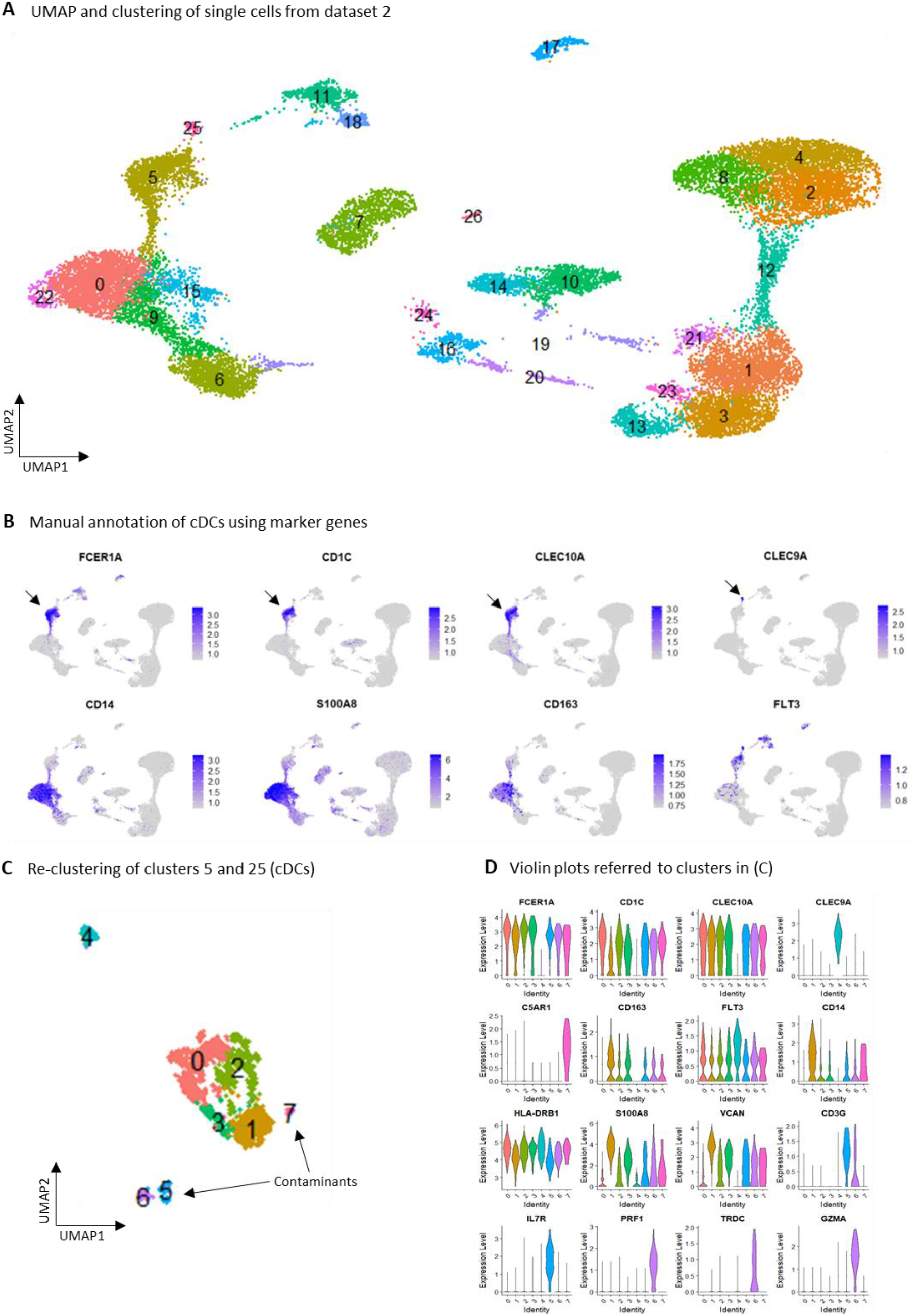
(A) UMAP and clustering of single cells from dataset 2. (B) Feature plots showing the expression levels of selected marker genes used to identify cDC clusters. Black arrows indicate cDC clusters (cluster 5 is cDC2 and cluster 25 is cDC1). (C) Re-clustering of clusters 5 and 25 corresponding to cDCs. Clusters 5, 6 and 7 were identified as contaminants. Clusters 0, 1, 2, 3 and 4 were re-clustered in a final iteration to clearly delineate cDC1, DC2 and DC3 subsets as shown in Figure 1B. (D) Violin plots referred to clusters in (C) showing expression levels of selected marker genes.

**Supplementary Figure 4.**
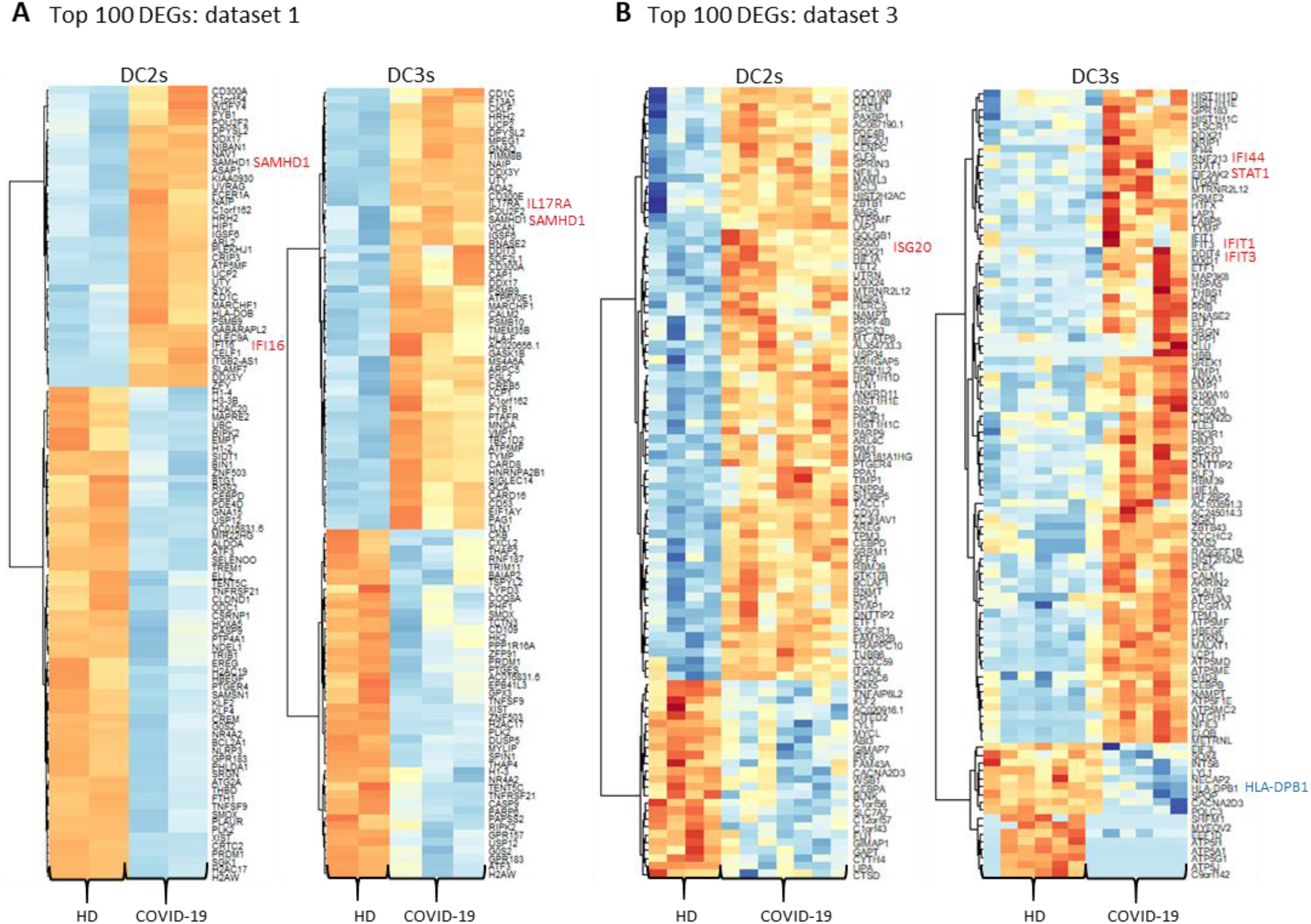
Heatmaps showing the top 100 DEGs for DC2 and DC3 subsets comparing COVID-19 patients and HDs from (A) dataset 1 and (B) dataset 2. Selected up-regulated genes are marked in red and down-regulated genes in blue. Ribosomal protein (RP) genes were removed from the top 100 DEGs.

**Supplementary Figure 5.**
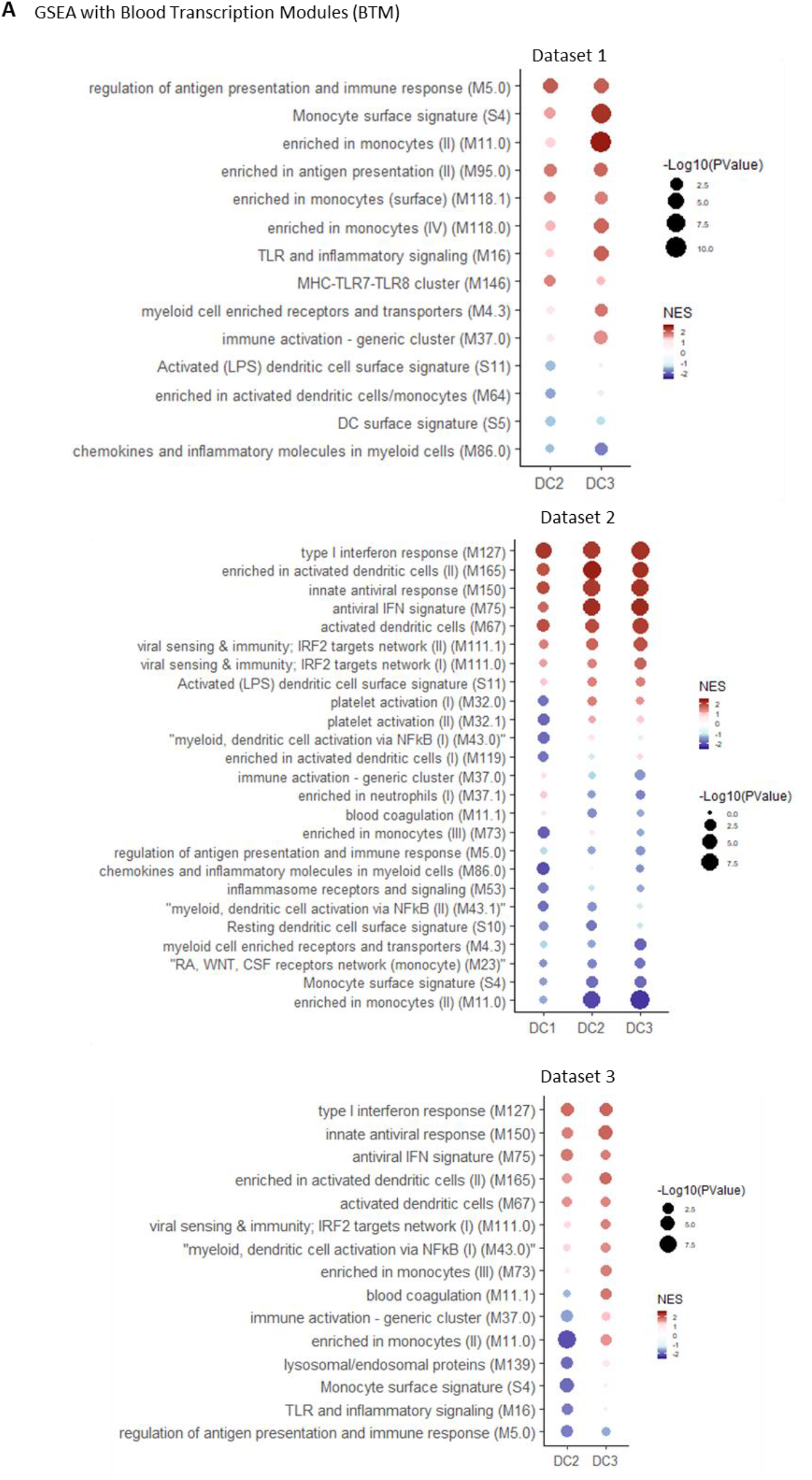
(A) GSEA of DEGs using the BTM collection: dataset 1 (upper panel), dataset 2 (middle panel) and dataset 3 (lower panel). For each DC subset, top 10 pathways based on significance are shown. NES, normalized enrichment score.

**Supplementary Figure 6.**
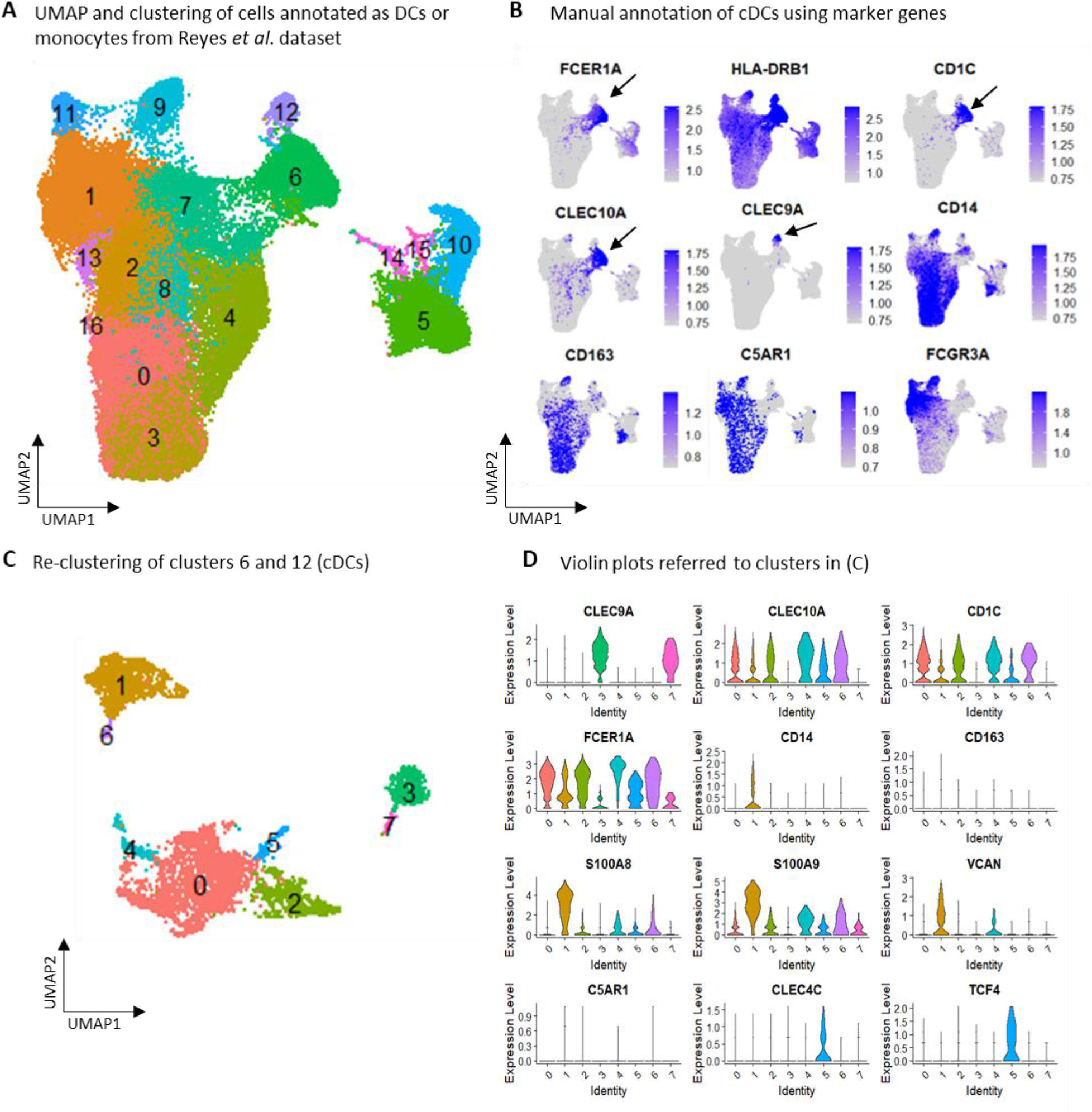
(A) UMAP and clustering of cells annotated as DCs or monocytes by the authors. (B) Feature plots showing the expression levels of selected marker genes used to identify cDC clusters. Black arrows indicate cDC clusters (cluster 6 is cDC2 and cluster 12 is cDC1). (C) Reclustering of clusters 6 and 12 corresponding to cDCs. (D) Violin plots referred to clusters in (C) showing expression levels of selected marker genes. Cluster 5, positive for CLEC4C and TCF4 was identified as contaminant and removed. All other clusters were re-clustered in a final iteration to clearly delineate cDC1, DC2 and DC3 subsets as shown in Figure 2C.

**Supplementary Figure 7.**
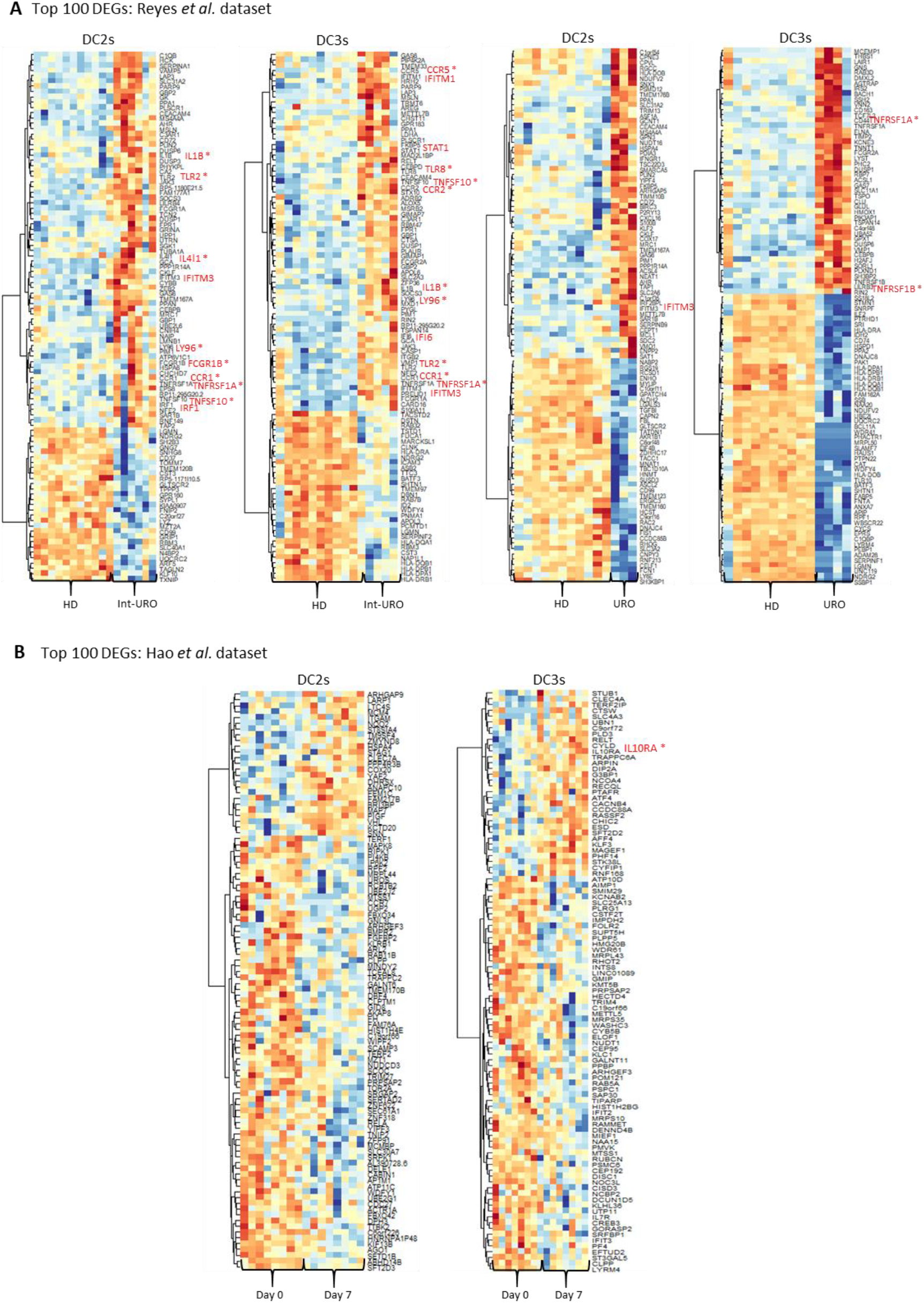
Heatmaps showing the top 100 DEGs for DC2 and DC3 subsets from (A) Reyes *et al*. and (B) Hao *et al*. datasets. Selected up-regulated genes are marked in red. Asterisk indicates genes associated with pro-inflammatory functions. Ribosomal protein (RP) genes were removed from the top 100 DEGs. Int-URO, intermediate urosepsis. URO, urosepsis.

**Supplementary Figure 8.**
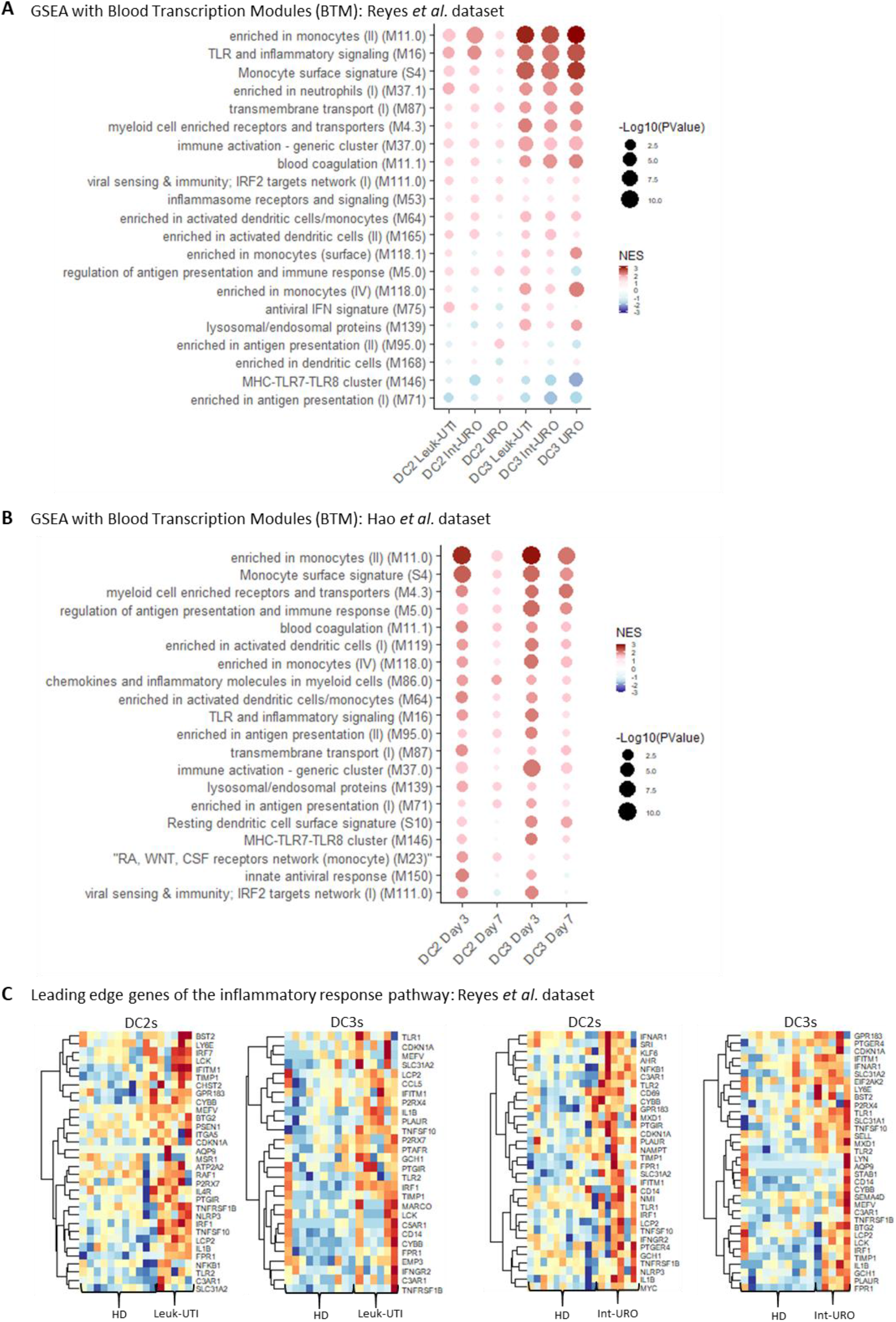
GSEA of DEGs using the BTM collection: (A) Reyes *et al*. dataset and (B) Hao *et al*. dataset. For each DC subset, top 10 pathways based on significance are shown. NES, normalized enrichment score. (C) Heatmaps showing leading edge genes of the inflammatory response pathway in the Reyes *et al*. dataset. Leuk-UTI, urinary tract infection with leukocytosis. Int-URO, intermediate urosepsis. URO, urosepsis.

**Table S1.** Characteristics of COVID-19 patients and healthy donors (HD) enrolled in the study.

| sample ID | Disease severity | age | gender | experiment |
| --- | --- | --- | --- | --- |
| Covid 1 | 5, SEVERE | 62 | female | scRNAseq |
| Covid 2 | 4, MILD | 66 | male | scRNAseq |
| Covid 3 | 4, MILD | 68 | female | scRNAseq |
| HD1 | N/A | N/A | N/A | scRNAseq |
| HD2 | N/A | N/A | N/A | scRNAseq |
| Covid 4 | 5, SEVERE | 87 | male | DCs frequency analysis |
| Covid 5 | 5, SEVERE | 68 | male | DCs frequency analysis |
| Covid 6 | 5, SEVERE | 49 | male | DCs frequency analysis |
| Covid 7 | 4, MILD | 79 | male | DCs frequency analysis |
| Covid 8 | 6, SEVERE | 77 | male | DCs frequency analysis |
| Covid 9 | 4, MILD | 60 | male | DCs frequency analysis |
| Covid 10 | 4, MILD | 81 | female | DCs frequency analysis |
| Covid 11 | 4, MILD | 84 | male | DCs frequency analysis |
| Covid 12 | 4, MILD | 67 | female | DCs frequency analysis |
| Covid 13 | 4, MILD | 69 | male | DCs frequency analysis |
| Covid 14 | 4, MILD | 89 | female | DCs frequency analysis |
| Covid 15 | 6, SEVERE | 59 | female | DCs frequency analysis |
| Covid 16 | 3, MILD | 53 | male | DCs frequency analysis |
| Covid 17 | 3, MILD | 89 | male | DCs frequency analysis |
| Covid 18 | 6, SEVERE | 61 | male | DCs frequency analysis |
| Covid 19 | 4, MILD | 80 | female | DCs frequency analysis |
| Covid 20 | 4, MILD | 79 | male | DCs frequency analysis |
| Covid 21 | 4, MILD | 85 | female | DCs frequency analysis |
| Covid 22 | 4, MILD | 59 | male | DCs frequency analysis |
| Covid 23 | 3, MILD | 72 | male | DCs frequency analysis |
| Covid 24 | 6, SEVERE | 62 | male | DCs frequency analysis |
| Covid 25 | 3, MILD | 78 | male | DCs frequency analysis |
| Covid 26 | 6, SEVERE | 68 | male | DCs frequency analysis |
| Covid 27 | 3, MILD | 67 | male | DCs frequency analysis |
| Covid 28 | 4, MILD | 83 | female | DCs frequency analysis |
| Covid 29 | 6, SEVERE | 31 | male | DCs frequency analysis |
| Covid 30 | 4, MILD | 79 | male | DCs frequency analysis |
| Covid 31 | 4, MILD | 80 | male | DCs frequency analysis |
| Covid 32 | 3, MILD | 72 | male | DCs frequency analysis |
| Covid 33 | 6, SEVERE | 65 | female | DCs frequency analysis |
| HD3 | N/A | 34 | female | DCs frequency analysis |
| HD4 | N/A | 32 | male | DCs frequency analysis |
| HD5 | N/A | 27 | male | DCs frequency analysis |
| HD6 | N/A | 40 | female | DCs frequency analysis |
| HD7 | N/A | N/A | N/A | DCs frequency analysis |
| HD8 | N/A | N/A | N/A | DCs frequency analysis |
| HD9 | N/A | N/A | N/A | DCs frequency analysis |
| HD10 | N/A | N/A | N/A | DCs frequency analysis |
| HD11 | N/A | N/A | N/A | DCs frequency analysis |
| HD12 | N/A | N/A | N/A | DCs frequency analysis |
| HD13 | N/A | N/A | N/A | DCs frequency analysis |
| HD14 | N/A | 57 | female | DCs frequency analysis |
| HD15 | N/A | 34 | male | DCs frequency analysis |
| HD16 | N/A | 41 | male | DCs frequency analysis |
| HD17 | N/A | 85 | male | DCs frequency analysis |
| HD18 | N/A | 62 | N/A | DCs frequency analysis |
| HD19 | N/A | 73 | N/A | DCs frequency analysis |
| HD20 | N/A | 57 | N/A | DCs frequency analysis |
| HD21 | N/A | 34 | N/A | DCs frequency analysis |
| HD22 | N/A | 70 | N/A | DCs frequency analysis |
| HD23 | N/A | 64 | N/A | DCs frequency analysis |

**Table S2. DEGs between COVID-19 patients and healthy donors in each DC subset from datasets 1, 2 and 3.**

**Table S3. BTM families used for GSEA.**

**Table S4. DEGs in each DC subset from Reyes *et al*. and Hao *et al*. datasets.**

**Table S5. DEGs in DC3s compared with DC2s in response to SARS-CoV-2 infection and intermediate urosepsis.**

**Table S6. DEGs in DC2s and DC3s in severe compared with mild patients.**

